# DNA supercoiling modulates bZIP transcription factor–DNA interaction

**DOI:** 10.64898/2026.05.06.722604

**Authors:** Rosanna Mattossovich, Rosa Merlo, Gioacchino Schifino, Annamaria Sandomenico, Mikael Widersten, Antonino Caliò, Judith Peters, Annalisa Pastore, Antonietta Parracino, Anna Valenti

**Affiliations:** Institute of Biosciences and BioResources, National Research Council of Italy, Via Pietro Castellino 111, 80131 Naples, Italy; Univ. Grenoble Alpes, CNRS, LiPhy, 140 rue de la physique St Martin d’Hères, 38400, France; Institute of Biostructures and Bioimaging, National Research Council of Italy, Via Pietro Castellino 111, 80131 Naples, Italy; Department of Chemistry-BMC, Uppsala University, BMC Box 576, Uppsala, S-751 23, Sweden; Institut Laue-Langevin, 71 avenue des Martyrs, CS 20156, 38042 Grenoble Cedex 9, France; Institut Universitaire de France, France; The Wohl Institute, King’s College London, 5 Cutcombe Rd, London SW59RT, U.K.; Elettra Sincrotrone Trieste, s.s. 14 km 163,500, Area Science Park, Basovizza, Trieste 34149, Italy

**Keywords:** DNA topology, leucine zippers, structure, DNA supercoiling, transcription factors, protein-DNA interaction

## Abstract

DNA topology is a key regulator of chromatin structure and transcription, yet its direct role in transcription factor recognition remains unclear. Here, we investigate how distinct DNA topological states modulate binding of the Saccharomyces cerevisiae bZIP transcription factor GCN4 using topologically defined plasmids. By combining, complementary biochemical approaches, including Bio-Layer Interferometry applied here for the first time to topology-dependent protein–DNA interactions, we show that DNA supercoiling directly reshapes GCN4–DNA recognition. Positively supercoiled DNA forms more stable and persistent complexes, whereas negatively supercoiled DNA retains greater conformational heterogeneity. To interpret these effects, we performed multiscale molecular simulations. Coarse-grained simulations of plasmids recapitulate the global topology-dependent trends observed experimentally, while matched minicircle models reproduce the same behaviour at the local scale. In strong agreement with experimental data, simulations reveal that DNA topology modulates the conformational ensemble of the GCN4 basic region. Overall, positively supercoiled DNA promotes a more ordered binding mode and localized protein distribution, whereas negatively supercoiled DNA supports increased structural plasticity. These findings identify DNA topology as an active determinant of transcription factor recognition and provide a multiscale framework linking global DNA mechanics to local protein–DNA interactions.

**Graphical Abstract:** 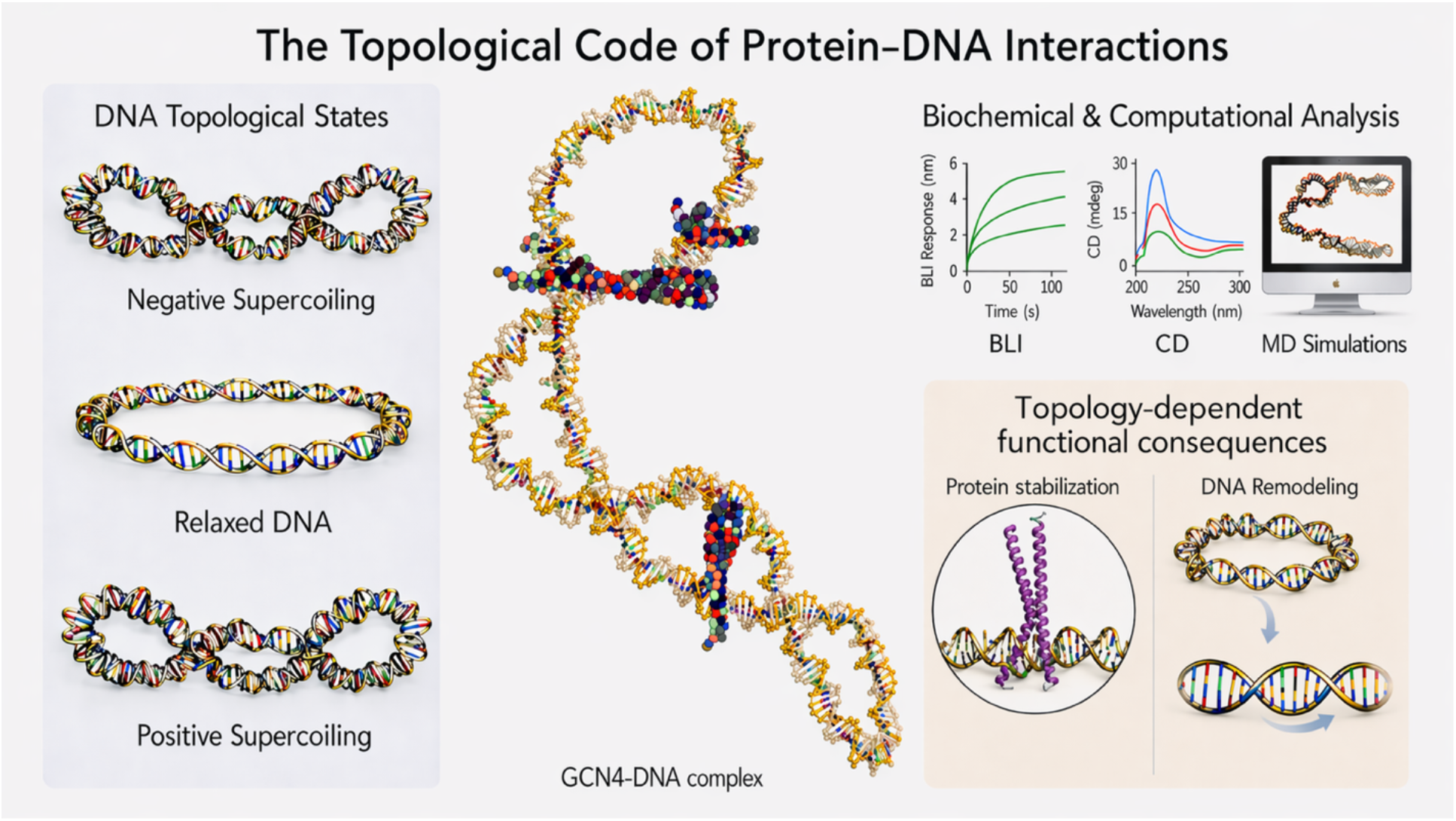

## 1. Introduction

While the iconic right-handed double helix is the most recognizable shape of DNA, in reality DNA rarely exists in this relaxed state inside a living cell. It is, instead, often either underwound or overwound, leading it to coil even further or “supercoil”. Recent discoveries have shed new light on the diverse roles of DNA supercoiling in chromatin regulation, while also highlighting the growing complexity of its functions (1-3). The double helix itself is a right-handed structure, but supercoiling introduces a higher-order chirality that can occur in two opposite senses, negative and positive, each with distinct structural and biological implications. Negative supercoiling is left-handed and thus occurs when the DNA is wound on the opposite handedness as compared to the double-helix chirality, thus resulting in underwinding of the chain with fewer twists than in its relaxed form. This makes the strands easier to separate, which is what is needed during processes like transcription or DNA replication, when the cell needs to “open” the helix. In bacteria, the entire genome is a single, circular piece of DNA that is usually, but not always, highly negatively supercoiled facilitating strand separation during DNA transactions (4,5). Conversely, positive supercoiling overwinds the DNA, tightening the helix and increasing its resistance to strand separation. This is quite helpful in certain contexts, such as in thermophilic organisms that live at high temperatures, where positive supercoiling stabilizes the DNA, preventing thermal denaturation. In eukaryotic cells, DNA is organized into chromosomes composed of topologically constrained domains and loops anchored by protein complexes. These constraints allow supercoiling to be generated and maintained locally, thereby modulating DNA accessibility and mechanical properties. Importantly, DNA topology is dynamically regulated: environmental cues, metabolic states, and cellular stress can induce rapid changes in supercoiling levels, providing a means to adjust the physical properties of the genome in response to changing conditions (6-8)

In this dynamic context, proteins emerge not only as passive responders but as active regulators of DNA topology. Non-covalent binding of transcription factors, chromatin-associated proteins and remodelers can induce local distortions in DNA conformation, transiently altering superhelical tension. In contrast, topoisomerases directly remodel DNA topology through covalent cleavage and re-ligation of the phosphodiester backbone, introducing or relaxing supercoils persistently. Together, these processes establish DNA supercoiling as a dynamic and regulated parameter of genome function (9-11).

The two chiral forms of supercoiling not only differ in energy and mechanical properties but also produce distinct higher-order DNA shapes and surface features, such as groove orientation, backbone spacing, and local helical tension. Experimental and theoretical studies have demonstrated that supercoiled DNA adopts distinct higher-order conformations with unique structural signatures. These features suggest that DNA topology can modulate protein binding beyond primary sequence recognition, contributing an additional layer of specificity through changes in DNA flexibility and surface topology (7,12). This topological regulation is particularly relevant in dynamic cellular processes such as transcription, replication, and stress response, where DNA undergoes continual changes in torsional strain (12-14).

Despite the recognized importance of DNA supercoiling in genome regulation, our understanding of how different chiralities and degrees of superhelical tension influence the recognition of DNA by binding proteins remains remarkably limited. While it is well established that DNA topology can facilitate or hinder protein binding through indirect structural effects, it is unclear to what extent proteins discriminate between negatively and positively supercoiled substrates or adapt their own conformations in response to distinct topological cues. Moreover, whereas DNA deformation induced by protein binding has been extensively studied, the reciprocal influence of DNA topology on protein conformational dynamics remains comparatively underexplored.

In this work, we address these questions by investigating how DNA topology modulates protein recognition using a well-defined biochemical model system. We focus on the interaction between plasmid DNA substrates with distinct supercoiling states and the yeast transcription factor GCN4. GCN4 is the prototypical example of a classic DNA-binding basic leucine zipper (bZIP) motif, conserved throughout yeast species, and required for amino acid biosynthesis in response to amino acid starvation (15). Although GCN4 is a yeast protein it serves as a model for mammalian bZIP transcription factors such as c-Jun, c-Fos, and ATF/CREB, whose dysregulation is linked to cancer, neurodegeneration, and inflammatory diseases (16,17). GCN4 binds to DNA as a parallel homodimer, recognizing and attaching to specific DNA sequences known as AP-1-like sites, typically 5’-TGACTC-3’ motifs or longer octa-nucleotide motifs, as well as related sequence variants that maintain the core TGA(G/C)T(C/A) consensus characteristic of AP-1–responsive elements. Independent studies have demonstrated that GCN4 binding is not solely determined by sequence-specific recognition but also by the local DNA conformation and chromatin environment (18). The bZIP domain includes a basic region that binds DNA and a leucine zipper that mediates dimerization. Upon DNA binding, the basic region forms α-helices that contact the major groove in a “scissor” arrangement. This conformational plasticity renders the bZIP domain particularly sensitive to changes in DNA shape and mechanical properties (19-21).

We combined biochemical and biophysical assays with coarse-grained and atomistic simulations to elucidate DNA structural conformations and to assess how defined DNA supercoiling modulates protein–DNA recognition. Using plasmid DNA as a model, we examined binding of the GCN4 bZIP domain by EMSA, circular dichroism (CD), and Bio-Layer Interferometry (BLI), enabling real-time quantification of topology-dependent interactions with supercoiled DNA. Our results demonstrate that DNA topology significantly affects protein-DNA complex formation, impacting binding affinity, structural organization and conformational dynamics. Together, these findings establish DNA supercoiling as an active determinant of protein recognition and provide a mechanistic framework for understanding how topological states contribute to the regulation of protein-DNA interactions.

## 2. Results and discussion

### 2.1 Production and characterization of DNA plasmids with different topologies

To assess the contribution of DNA topology to protein-DNA interactions, plasmid DNA (pDNA) was prepared in four defined topological states: negatively supercoiled pDNA(SC-), open-circular pDNA(OC), linear pDNA(LIN), and positively supercoiled pDNA(SC+) (Figure 1A). These conformations display structural and mechanical differences expected to influence protein binding (7,22). The degree of supercoiling in covalently closed DNA is expressed by the superhelical density (σ), defined as σ= ΔLk/Lk_0_, where ΔLk is the change in linking number relative to the relaxed form (Lk_0_).

**Figure 1.**
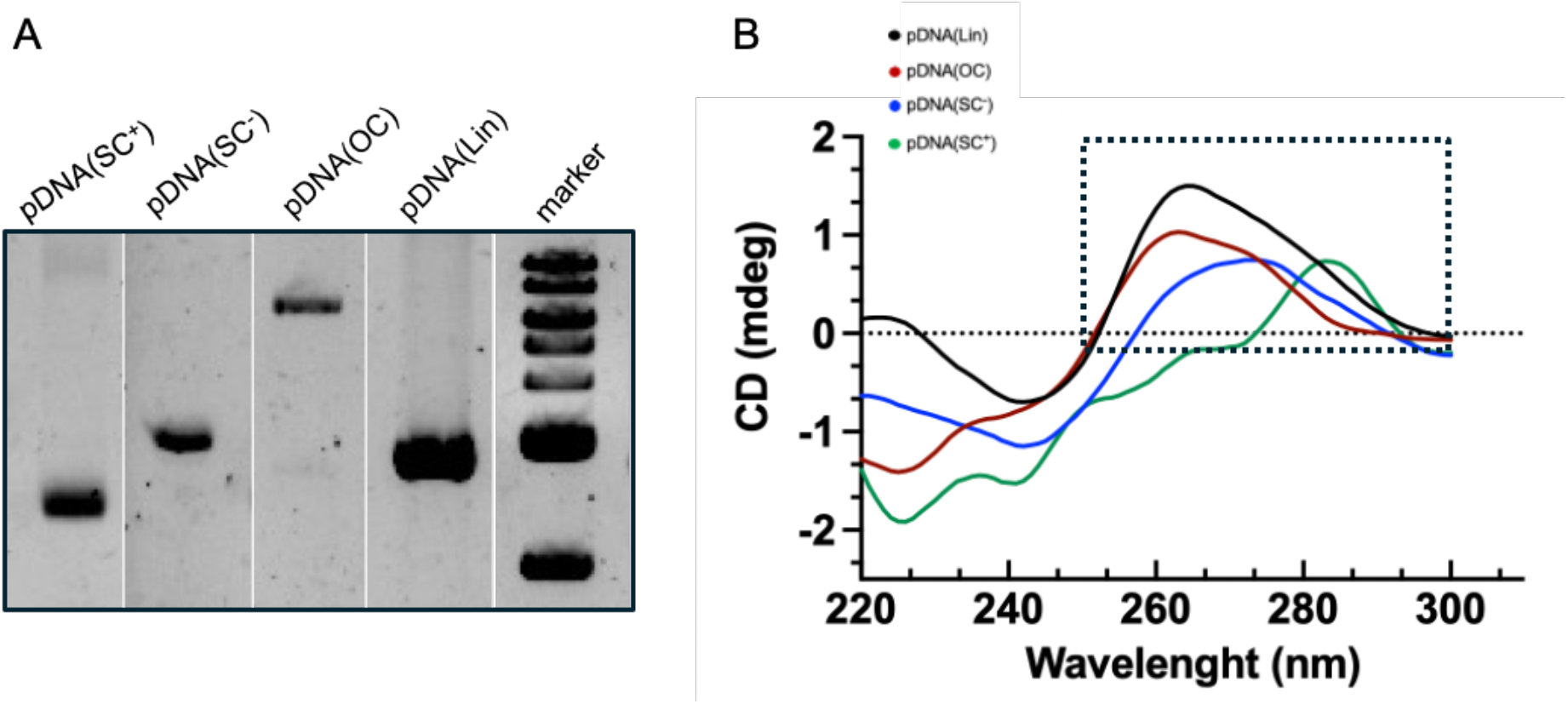
A) Topological separation of plasmid DNA (pDNA) by agarose gel electrophoresis. Samples were run on a 1.2% agarose gel Tris-Borate-EDTA (TBE) buffer. Lane 1: pDNA(SC+); lane 2: pDNA(SC-); lane 3: pDNA(OC); lane 4: pDNA(LIN); lane 5: 1 kb DNA ladder (range: 0.5-10 kb). B) Far- and near-UV CD spectra of pDNA in four topological states. Spectra (200-300 nm) were recorded at 25 °C with a 0.1 cm path-length cuvette using 16nM pDNA. Curves are colour-coded by topology: pDNA(LIN) in black, pDNA(OC) in red, pDNA(SC-) in blue and pDNA(SC+) in green. The rectangle highlights the near-UV window (260-300 nm) where topology-dependent shifts are most pronounced: pDNA(LIN) and pDNA(OC) peak at ∼265 nm, pDNA(SC-) red-shifts to ∼270 nm with reduced ellipticity, and pDNA(SC+) shifts further to ∼285 nm with additional amplitude loss. Each curve represents the average of three scans and is plotted as raw ellipticity (mdeg).

The pUC19 plasmid (2686 bp) was used as a well-established model substrate. Plasmid DNA isolated from *Escherichia coli* typically carries a negative σ of ∼-0.04 owing to DNA gyrase activity; this native form was used as pDNA(SC-). The open-circular form pDNA(OC) (α= 0) was generated by site-specific nicking, whereas linear pDNA(LIN) was obtained by restriction digestion with EcoRI. Positively supercoiled pDNA(SC+) (σ > +0.04) was produced using a thermophilic *reverse gyrase*, the only known enzyme able to introduce positive supercoils into DNA molecules. This workflow enabled generation of pDNA substrates sharing identical sequence and length while differing exclusively in topological state (7,23,24). To define the conformational signatures associated with each topology, we analyzed the resulting plasmids by CD spectroscopy. CD is a sensitive probe of chiral secondary structure and base stacking in nucleic acids, and classic studies showed that DNA with different topologies exhibit distinct near-UV CD signatures consistent with topology-dependent perturbations in base stacking and helical geometry (25,26).

Consistent with previous studies, CD spectroscopy showed that each pDNA displayed the canonical B-DNA profile with a dominant positive band at ∼275 nm, a shoulder near 220 nm and a negative band at ∼245 nm (Figure 1B). Topology-dependent shifts were also evident in the near-UV region (250-300 nm), which is sensitive to local changes in DNA base arrangement and stacking. Beyond this shared B-DNA framework, clear topology-dependent differences emerged in the 250-300 nm window: pDNA(OC) and pDNA(LIN) showed a single maximum at ∼265 nm, whereas pDNA(SC-) exhibited a red shift to ∼270 nm with reduced ellipticity, and pDNA(SC+) showed a further red shift to ∼285 nm with an additional loss of intensity. These progressive red shifts and intensity changes are consistent with prior reports linking superhelical stress to local alterations in base stacking and groove geometry and demonstrate that supercoiling imposes distinct conformational signatures readily detectable in the near-UV CD spectrum, adding structure-encoded complexity beyond the canonical B-DNA profile (22,27,28).

By directly comparing the same plasmid sequence across pDNA(SC-), pDNA(OC), pDNA(LIN), and pDNA(SC+), our analysis defines topology-specific CD signatures of DNA and establishes a reference for detecting and interpreting protein-induced conformational changes under defined supercoiling.

### 2.2 Topology-Dependent Gcn4–DNA Interactions Characterized by EMSA and an implemented BLI approach

To gain an initial view of how DNA topology influences GCN4 binding, we used EMSA, a classical method for detecting DNA–protein complexes under native conditions.

EMSA demonstrated that GCN4 associates with all DNA topologies tested, as indicated by the formation of shifted bands in each condition (Figure 2). Notably, the electrophoretic mobility of the protein-DNA complexes depended on the topology of the starting plasmid, consistent with topology-dependent differences in complex conformation. Similar mobility patterns were observed when the same plasmid contained a canonical GCN4 consensus site, suggesting that, in the context of large DNA substrates, global DNA topology plays a predominant role over local sequence determinants in GCN4-DNA interaction (data not shown). Although EMSA allowed qualitative discrimination among complexes, quantitative parameters were estimated using a recently published densitometric approach (29), yielding a dissociation constant (k_d_) that suggested higher affinity for supercoiled forms.

**Figure 2.**
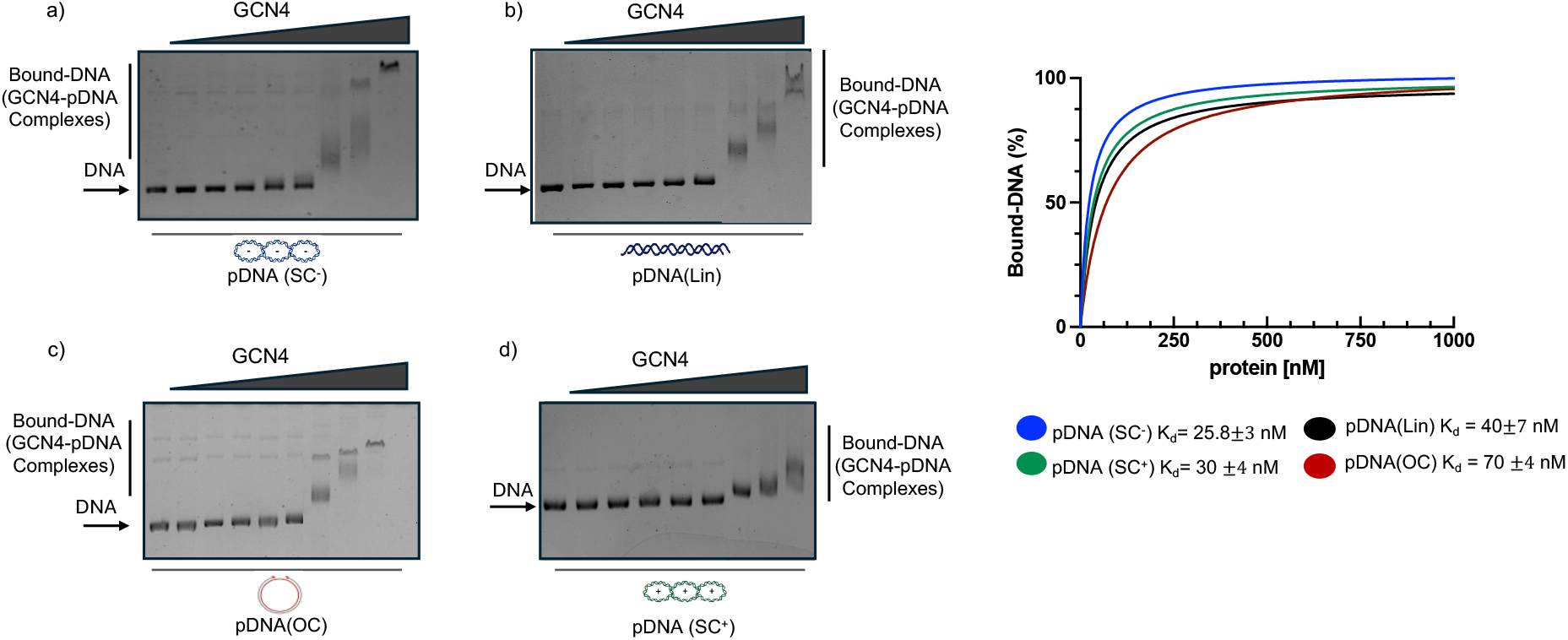
Analysis of the GCN4-pDNA interaction by EMSA. A) Increasing concentrations of GCN4 (5 to 1000 nM) were incubated for 10 minutes at 37 °C in the presence of pDNA (3 nM) of the indicated topologies: a) pDNA(SC-); b) pDNA(LIN); c) pDNA(OC); d) pDNA(SC+). DNA-protein complexes were analysed on a 1.2% agarose gel and visualized after ethidium bromide staining. The fraction of bound pDNA (GCN4-pDNA complexes) in lanes 2-9 was quantified relative to the sample containing no protein (lane 1) for each DNA topology. Data were fitted using a one-site binding equation to determine the dissociation constant (K_d_). The panel on the right shows representative binding curves corresponding to the EMSA datasets.

However, this analysis is intrinsically limited, as it does not account for topology-specific effects on DNA mobility, intercalating dye binding, or charge distribution, all of which may influence band intensity and migration. These considerations motivated the implementation of BLI, which we optimized here to investigate GCN4 interactions with plasmid DNA substrates of defined supercoiling.

Although BLI is widely used to characterize protein-protein and protein-antibody/small-molecule interactions, its application to DNA-protein systems remains relatively limited (30,31). In the BLI assay, biotinylated double-stranded oligonucleotides bearing a high-affinity GCN4 recognition site were immobilized on streptavidin biosensors to establish a reference interaction. Association and dissociation curves collected across a range of protein concentrations enabled estimation of kinetic parameters and identification of the optimal protein concentration for subsequent experiments (Fig. S1). This configuration allowed us to implement a competition-based BLI (cb-BLI) assay to compare the ability of plasmids with distinct topologies to displace GCN4 from the immobilized oligonucleotide (Figure 3A) (32). During the dissociation phase, pDNA was introduced as a competing ligand to promote protein displacement from the oligonucleotide in favor of pDNA. This indirect competition strategy overcomes technical limitations associated with direct plasmid immobilization, such as constrained orientation, steric hindrance and reduced conformational freedom, providing a more faithful representation of binding to topologically distinct substrates.

**Figure 3.**
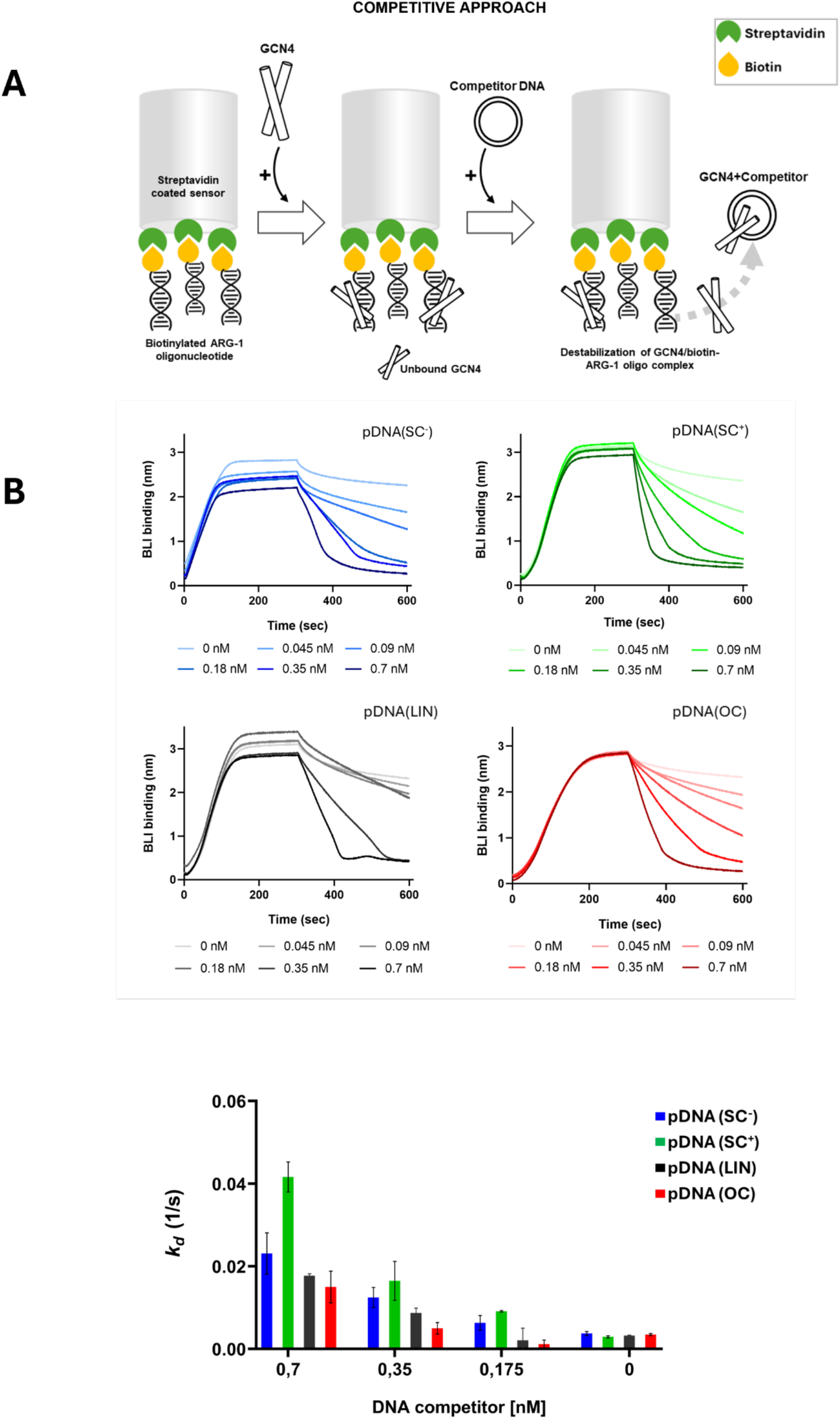
Competition experiments to study DNA-protein interaction by BLI. (A) General scheme for studying GCN4/DNA interaction using streptavidin-coated sensors on BLI. After loading fixed concentrations of the GCN4 protein onto bio-oligo-coated biosensors, the *k*_*d*_ was measured in the presence of different concentrations of plasmid DNA with different topologies, used as competitors. The measured dissociation curves reflect the affinity of GCN4 for the plasmid DNA and the consequent destabilisation of the immobilised protein/oligo complex; (B) Association and dissociation curves of a fixed concentration of GCN4 (0.1 µM) in the presence of various concentrations of DNA competitors for all the topologies analysed (pDNA(SC-) in blue, pDNA(SC+) in green, pDNA(LIN) in grey and pDNA(OC) in red). The dissociation curves and dissociation constants were analysed after immersing the pre-formed GCN4/oligo complex in wells containing DNA with different topologies, as potential competitors. Bar chart representation of the topology-dependent increasing *k*_*d*_ and the relative affinity of GCN4 for plasmid DNA based on the topology.

The presence of competing pDNA resulted in an increased dissociation rate of GCN4. The recorded dissociation rates were compared to the *k*_*d*_ values obtained from using pure buffer and correlated with the relative affinity of the protein for a given pDNA topology, with faster dissociation indicating stronger binding to the competitor (Figure 3B). Interestingly, comparison of the *k*_*d*_ showed an increase of GCN4 displacement in the presence of supercoiled pDNA as compared to the linear and open circular pDNA forms, with the strongest binding observed for the pDNA(SC+).

The binding preferences observed in the cb-BLI assay can be rationalized by considering the distinct biophysical features of the different DNA supercoiling states. Negatively supercoiled pDNA(SC-) stores torsional energy that promotes local unwinding, base flipping and transient denaturation, thereby increasing exposure of the phosphate backbone and enhancing electrostatic engagement with the positively charged basic region of GCN4. This provides a straightforward explanation for the higher affinity of GCN4 for pDNA(SC-) compared with pDNA(OC) and pDNA(LIN) substrates (33). In contrast, positively supercoiled pDNA(SC+) is more compact and less prone to strand separation, yet overwinding can induce localized distortions in helix geometry, altering groove dimensions and generating sterically favorable binding pockets. Such features may facilitate tighter docking of the bZIP dimer and reduce its dissociation rate, offering a plausible structural basis for the strong binding observed for pDNA(SC+). The pronounced displacement of GCN4 toward pDNA(SC+) therefore, suggests that overwound DNA can stabilize the complex not only through electrostatic complementarity but also via topological confinement. Together, these data support a model in which DNA supercoiling modulates both the accessibility and the conformational landscape of GCN4-DNA complexes, with pDNA(SC+) providing the most favorable binding geometry under our experimental conditions. Given the dynamic nature of local DNA supercoiling in vivo, our findings suggest that positive supercoiling could influence transcription factor retention or preferential recruitment at specific genomic loci (34).

### 2.3 Structural Insights into Gcn4–DNA Complexes by Circular Dichroism

We next used far-UV CD to evaluate whether DNA topology influences the structural properties of the GCN4–DNA complexes. This is a difficult territory because, while this technique is uniquely sensitive to chirality and would thus be very appropriate for these studies, it is also not able to discriminate among the individual contributions in a complex. Any changes in the far-UV CD spectra of the complex will report both on the protein and on the plasmid in a non-linear way. We thus used this technique carefully. The CD spectrum of GCN4 in solution shows the expected α-helical signature, with minima at 208 and 222 nm and a maximum near 190 nm, consistent with previous reports (35,36).

Upon titration with plasmid DNA, we observed pronounced alterations in the protein–DNA complex spectra: ellipticity increased at ∼208 nm, whereas the canonical 222 nm minimum shifted toward ∼230 nm with reduced intensity. Because these signals arise from both components, they cannot be interpreted solely in terms of conventional protein secondary structure, but they clearly indicate that binding induces a conformational reorganization involving both GCN4 and plasmid (Figure 4). Interestingly, the most substantial changes occurred with positively supercoiled pDNA(SC+), whose complex spectrum exhibits a prominent minimum near 230 nm, a feature reminiscent of β-type signals but here more likely reflecting mixed contributions from altered DNA geometry and protein rearrangement. Similar far-UV profiles have been previously associated with high-affinity GCN4–DNA complexes on short duplexes, consistent with tight binding (36,37).

**Figure 4.**
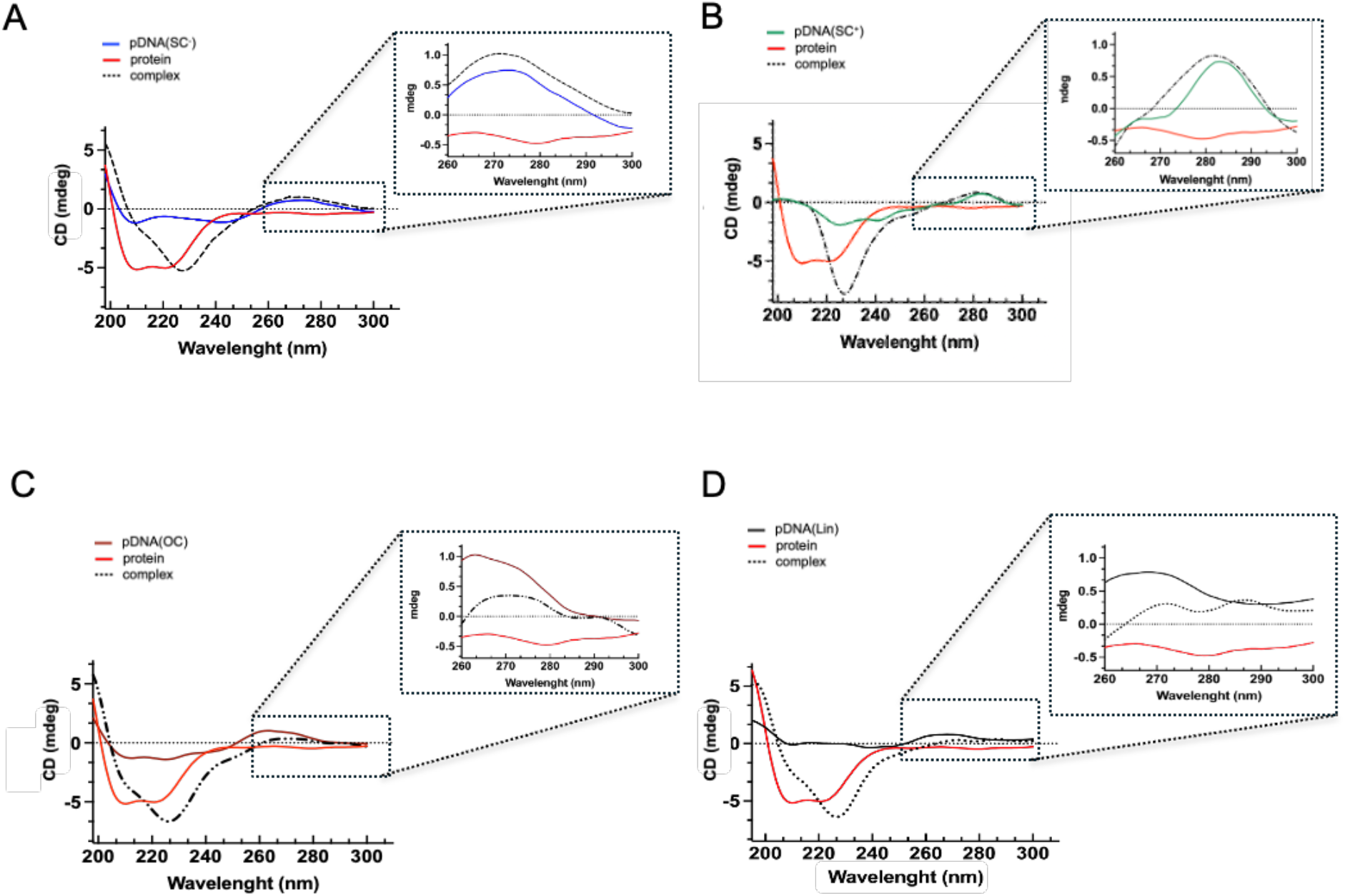
CD spectroscopy of GCN4 folding and plasmid DNA topology in protein-DNA interactions. Spectra were recorded at 25 °C using a 0.1 cm pathlength quartz cuvette, scanning from 200 to 300 nm (average of three accumulations). Each panel (A–D) displays the CD spectra of pDNA (16 nM) of a specific topology, either free (colored trace) or in complex with GCN4 (5 μM, black trace). The CD spectrum of GCN4 protein alone (5 μM) is shown in red in all panels. Free DNA topologies are color-coded as follows: (A) pDNA(SC-) in blue; (B) pDNA(SC+) in green; (C) pDNA(OC) in dark red; (D) pDNA(LIN) in black. GCN4–DNA complex spectra are shown in black. Insets highlight the near-UV region (260–300 nm), where GCN4 binding induces topology-dependent spectral changes. pDNA(SC-) exhibits a red shift and reduced ellipticity upon complex formation, while pDNA(SC+) shows a modest increase in ellipticity with minimal peak displacement. pDNA(LIN) and pDNA(OC) forms display intermediate behaviors.

To specifically analyze DNA structural changes, we focused on the 260-300 nm region, where the CD signal is dominated by plasmid contributions and minimally affected by the protein. Upon protein binding, a significant shift in the pDNA CD bands was observed for all topologies tested, indicating conformational changes in DNA structure after protein interaction. For pDNA(OC) and pDNA(LIN), binding caused a general decrease in CD intensity accompanied by a red shift, consistent with local alterations in base stacking. In contrast, the supercoiled substrates exhibited a different response: the CD signal slightly increased and the spectrum became more structured, indicating that DNA in the complex is more constrained. This suggests that the intrinsic torsional stress of supercoiled DNA limits conformational changes upon protein binding. The effect was less pronounced for pDNA(SC+) than for pDNA(SC-), consistent with the higher torsional rigidity imparted by overwinding, which may restrict DNA flexibility and influence the docking geometry of the GCN4 dimer. Taken together, these results indicate that GCN4 not only senses DNA topology but also induces topology-dependent structural adjustments in DNA. Conversely, DNA supercoiling modulates the conformational landscape available to the protein. This bidirectional structural coupling supports a model in which the architecture of the GCN4-DNA complex is jointly shaped by both partners, with positive supercoiling promoting the most pronounced and constrained rearrangements.

### 2.4 Multiscale Analysis of Conformational Dynamics in circular DNA systems

To place the local observations described above in a broader structural context, we next examined the topology-dependent conformational behavior of the circular DNA within a multiscale framework. Although positive DNA supercoiling exhibits distinct structural and biophysical properties that, as demonstrated by the experimental work reported here, influence protein/DNA interactions, it remains far less explored than negative supercoiling, particularly in multiscale simulations of long plasmid substrates (5,7,8,22,26). Coarse-grained analyses of circular DNA systems spanning plasmid-sized constructs and smaller torsionally stressed minicircle have shown that superhelical stress can strongly reshape the accessible conformational ensemble, yet positively supercoiled systems remain comparatively underrepresented, especially in frameworks that directly connect plasmid-scale behavior to matched local models (38-40). For this reason, we used coarse-grained simulations of the plasmid to define the global topology-dependent conformational bias of the experimentally interrogated substrate.

At the plasmid scale, the trajectories define a clear preference trend. pDNA(SC+) occupies a narrower and more compact conformational basin, whereas pDNA(SC-) samples a broader and more heterogeneous landscape (Figure 5, while a snapshot of -pDNA(SC+) is in supplementary Figure S12). This behavior is important for two reasons: first, it highlights that the positively supercoiled plasmid is not simply an alternative isomeric state, but a structurally distinctive conformational regime; second, it provides the global topological reference needed to interpret the atomistic mcDNA simulations in the correct mechanical context. In this sense, the plasmid course-grain analysis captures a topological state that remains comparatively less explored in plasmid-scale multiscale simulations and establishes the large-scale structural bias from which the local observations can be read.

**Figure 5.**
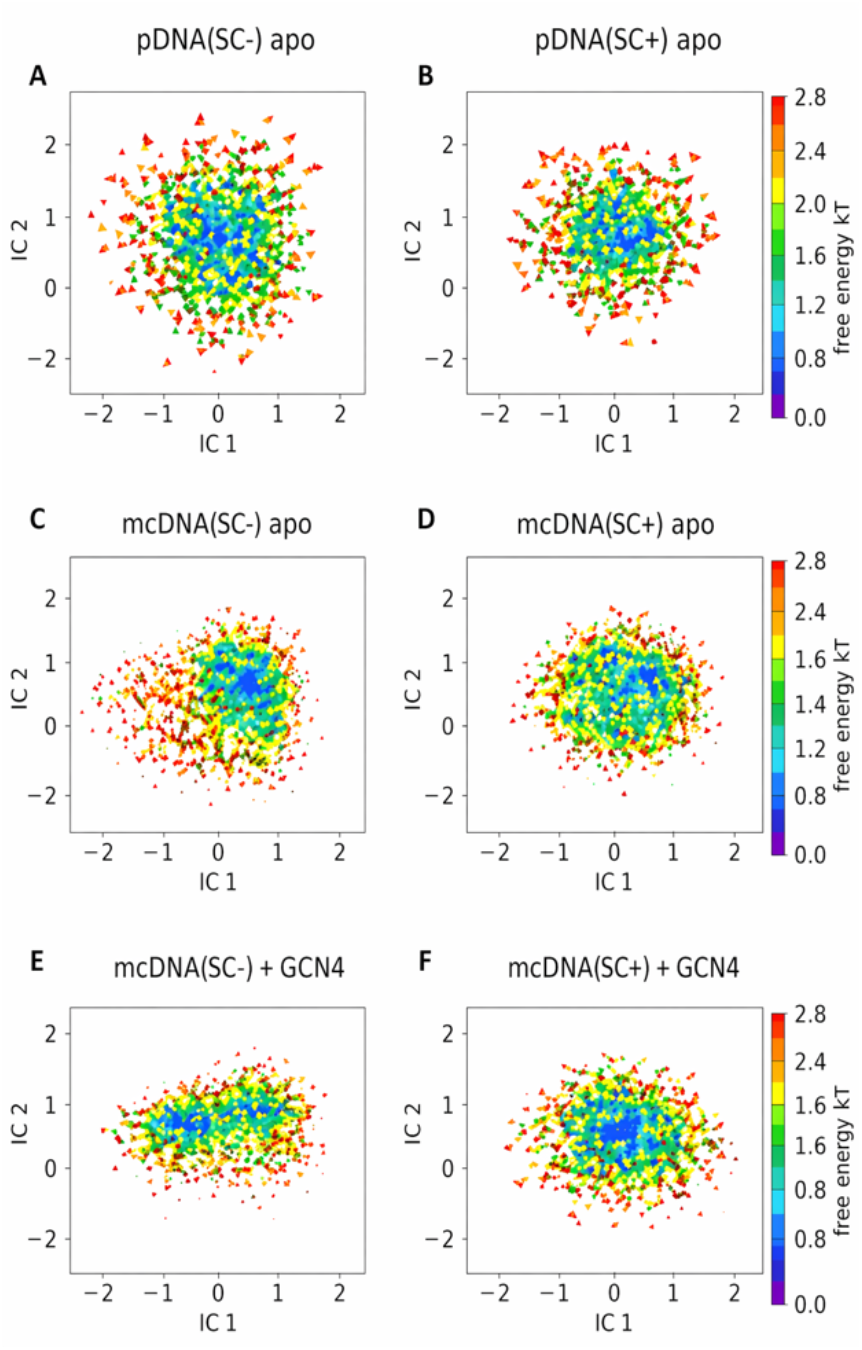
Integrated multiscale conformational landscapes. Panels A-B show the plasmid coarse-grained tICA/FES projections for pDNA(SC-) and pDNA(SC+), panels C-D show the corresponding apo mcDNA landscapes, and panels E-F show the GCN4-bound mcDNA landscapes. The figure is used to establish cross-scale consistency of the topology-dependent landscape, not absolute equivalence between models.

To gain atomistic-level insight, we simplified the system by adopting a 336-bp DNA minicircle (mcDNA), a reduced circular construct which preserves the topological behaviour of longer plasmids while significantly reducing computational resources. The corresponding coarse-grained plasmid segment was extracted frame by frame from the plasmid trajectories and compared with the matched apo mcDNA systems. Across these contexts, the negative state retained a larger radius of gyration than the positive state, indicating that the compaction bias observed at the plasmid scale is preserved locally (Supplementary Figure S6). Importantly, this agreement should be interpreted at the level of topology-dependent ordering rather than absolute structural equivalence, since coarse-grained and atomistic models are not expected to yield identical numerical results. This cross-scale agreement is the key rationale for using mcDNA as a mechanistically informative local proxy rather than as an substitute for the whole plasmid. Together, the free-energy results and the radius-of-gyration reported in Figure 5 and Supplementary Figure S6 define a coherent multiscale argument: the plasmid establishes the global topology-dependent bias, while the matched local systems preserve the same sign-dependent ordering at the scale used for atomistic analysis. The mcDNA models can therefore be interpreted as locally resolved expressions of a broader plasmid response, providing a direct conceptual bridge between large-scale DNA mechanics and the protein-coupled structural behavior described in the previous section.

### 2.5 Minicircle DNA Models to Probe the Dynamic Interplay Between DNA Topology, Protein Conformation, and Binding Energetics

To probe how DNA topology modulates the local organization of the complex, we used protein-bound mcDNA systems as structurally resolved models of the GCN4-DNA interaction. Representative snapshots of mcDNA(SC−), mcDNA(SC+), and of the corresponding GCN4-bound complexes are shown in Supplementary Figures S2 and S3, respectively. In this context, the minicircle is not treated simply as a reduced DNA substrate, but as a local framework in which the consequences of topological stress can be examined at a level of detail that is not directly accessible in the plasmid. Because all mcDNA systems share the same sequence framework while differing in topological state, they provide a suitable setting to evaluate how local DNA geometry and torsional stress influence protein conformational dynamics, local DNA rearrangement, and binding-related structural descriptors around the specific site.

The clearest topology-dependent signal emerged from the conformational plasticity of the bZIP basic region. While the leucine-zipper dimerization core remained comparatively stable in all systems, the DNA-binding arms displayed a markedly different behavior depending on topology. In mcDNA(SC-), the basic region sampled broader rearrangements and a wider conformational space, whereas in mcDNA(SC+) it was confined to a more ordered and restricted ensemble (Supplementary Figure S5). Residue-resolved secondary-structure persistence maps and per-dimer RMSF/helicity profiles further support this interpretation, showing that the positively supercoiled systems retain a more persistent helical organization within the DNA-binding region, whereas mcDNA(SC-) displays broader local fluctuations across the same structural elements (Supplementary Figures S8 and S9). This distinction is mechanistically informative because it indicates that DNA topology does not merely influence the probability of binding, but also reshapes the accessible conformational landscape of the protein once bound. The more ordered ensemble observed on mcDNA(SC+) is also consistent with the tighter and more persistent complexes inferred from the experimental measurements.

Local DNA readouts followed the same general logic. Protein-DNA hydrogen-bond persistence and the local DNA radius of gyration did not collapse into a single absolute ranking, but instead redistributed in a topology-dependent manner across the four atomistic mcDNA systems (Supplementary Figure S7). We interpret these descriptors as local manifestations of how each topological state reorganizes the protein-DNA interface. In this view, the immediate binding environment reflects a balance between DNA torsional stress, local curvature, and the ability of the protein to adopt a productive binding geometry. A further level of organization emerged from the positional analysis of GCN4 dimers along the minicircle, where five dimers were initially positioned on the same mcDNA template. In all systems, the dimer docked on the specific AP-1 site remained the persistent primary partner, whereas topology mainly redistributed the remaining non-primary dimers around the same mcDNA template. This distinction is important because it separates chemically encoded primary recognition from topology-dependent secondary organization. In mcDNA(SC+), one non-primary dimer recurrently emerged as a dominant secondary partner, whereas mcDNA(LIN), mcDNA(NICK), and mcDNA(SC-) retained a broader and less focused occupation pattern (Figure 6).

**Figure 6.**
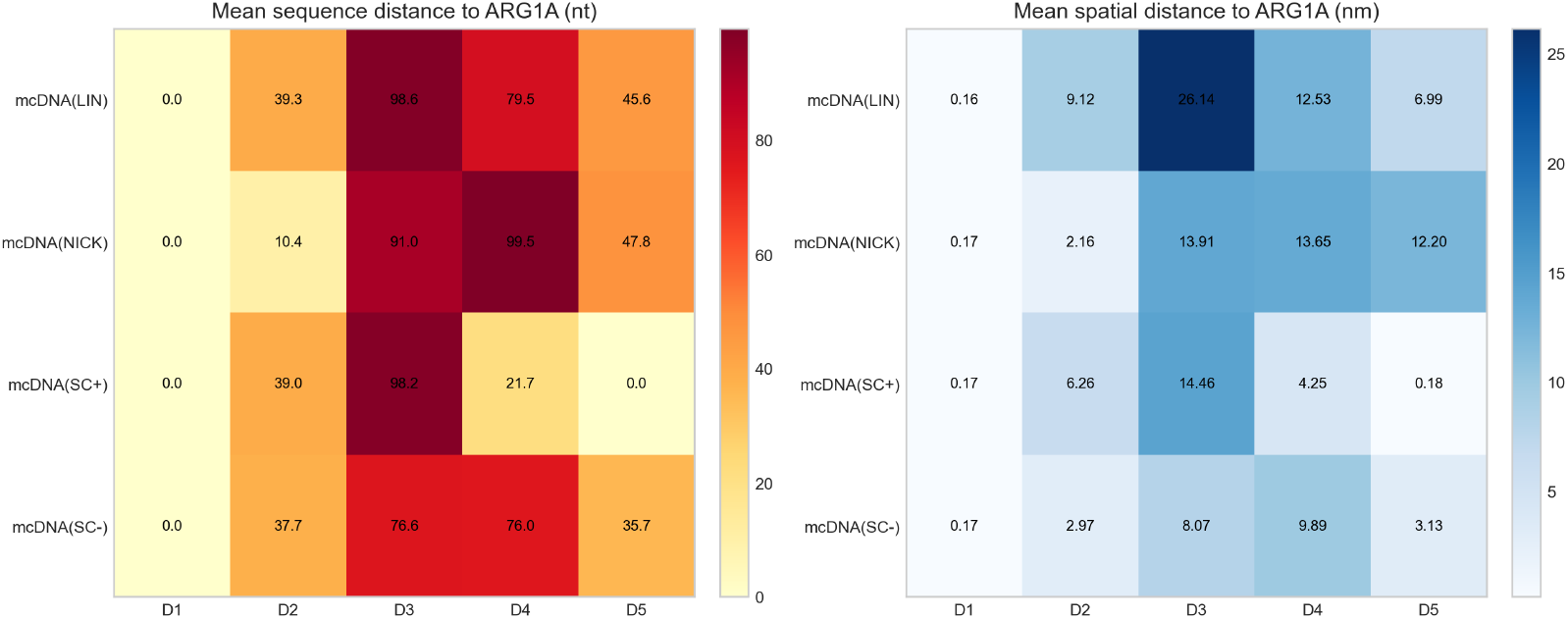
Heatmap analysis of dimer placement relative to the AP-1 specific site across the four mcDNA systems. The left panel reports the mean sequence distance (nt) of each dimer from the AP-1 specific site, whereas the right panel reports the corresponding mean spatial distance (nm). Together, these descriptors resolve the hierarchy between the persistent primary dimer and the topology-dependent redistribution of the non-primary dimers.

The same trend is reinforced when the best non-primary approach to canonical AP-1 and AP-1-like motifs is ranked explicitly and when the occupancy of motif-compatible landing sites is visualized across the minicircle (Figure 7 and in Supplementary Figure S10).

**Figure 7.**
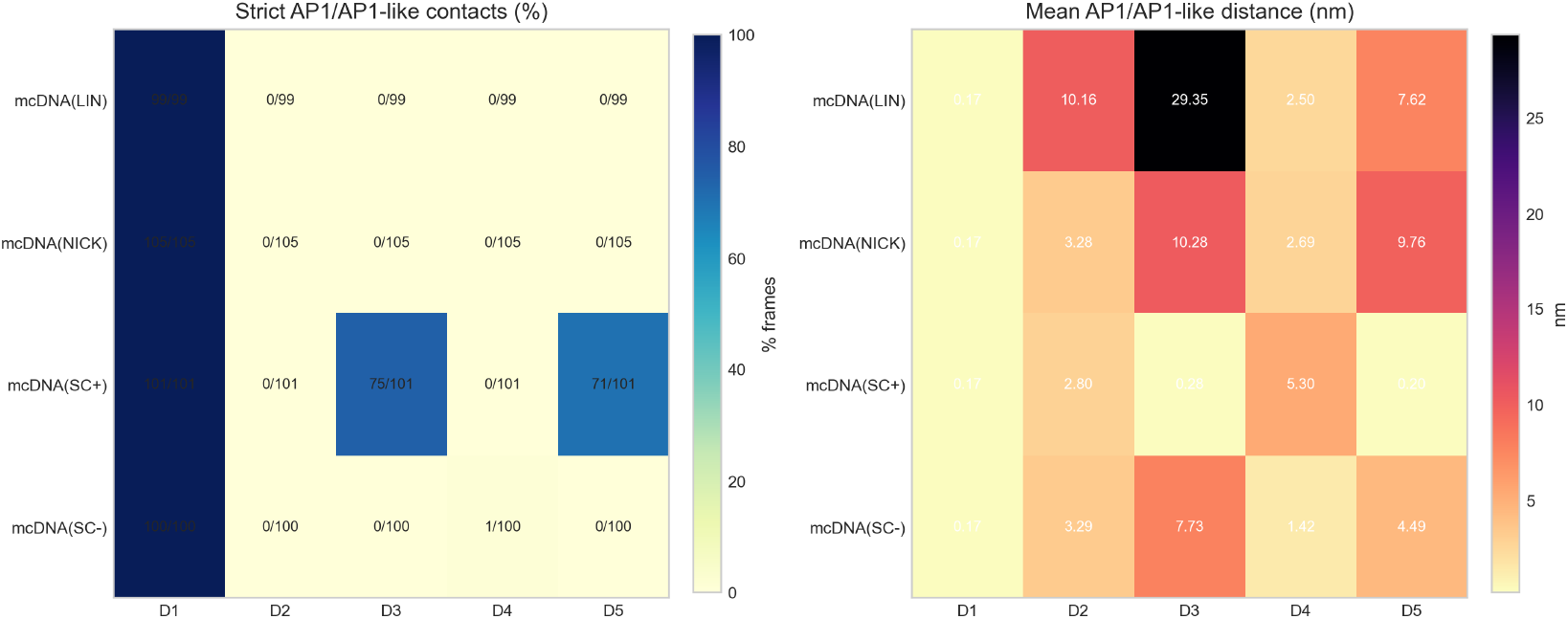
Motif-aware analysis of non-primary dimer positioning across the mcDNA systems. The left panel summarizes the best AP-1/AP-1-like contacts identified for the non-primary dimers, whereas the right panel reports the corresponding mean approach distances, providing a complementary view of topology-dependent redistribution toward motif-compatible regions.

Comparative MM/GBSA analyses, retained only in the SI as supportive energetic descriptors, are consistent with this interpretation and support the within-study ranking of the mcDNA systems (Supplementary Figure S11). Taken together, these results support a two-level model in which sequence-specific docking establishes the primary anchoring event, whereas DNA topology modulates how additional dimers are redistributed across alternative DNA segments and how the local conformational ensemble of the complex is stabilized.

## 3. Conclusions

Our results show that DNA topology is not a passive background feature of the substrate, but an active determinant of GCN4–DNA recognition. By combining EMSA, competition Bio-Layer Interferometry, circular dichroism, and multiscale simulations, we demonstrate that distinct topological states of the same plasmid sequence give rise to measurably different binding behaviors and structural responses. In particular, positively supercoiled pDNA supports stronger and more persistent complexes, whereas negatively supercoiled pDNA is associated with broader conformational heterogeneity. These observations indicate that the topological state of DNA contributes directly to shaping both the stability and the structural organization of the resulting protein–DNA complex.

In this work, we take advantage of topologically defined plasmids as experimentally tractable substrates to isolate the effects of DNA topology. Distinct topological states can be generated in a controlled manner through enzymatic treatments, allowing access to plasmids with defined supercoiling levels and signs. Because they are covalently closed, these molecules preserve the coupling between twist and writhe, thereby maintaining the mechanical constraints that characterize supercoiled DNA *in vivo*. Within this context, our competition-BLI strategy provided a particularly informative way to compare relative binding preferences without imposing additional topological constraints through direct plasmid immobilization. Together with the CD measurements, these experiments show that GCN4 does not simply bind DNA sequence in an invariant manner, but responds to the structural and mechanical properties encoded by DNA topology. The CD data further support this interpretation by revealing topology-dependent chiroptical signatures of the complexes, consistent with binding-induced rearrangements that depend on the initial topological state of the plasmid.

The computational analysis provides a mechanistic framework for interpreting these observations across scales. At the local level, atomistic simulations of protein-bound mcDNA systems show that DNA topology strongly affects the conformational plasticity of the GCN4 basic region, whereas the leucine-zipper core remains comparatively stable. In addition the analysis support a conceptual separation between sequence-encoded recognition and topology-dependent organization of protein–DNA complexes: while the specific DNA sequence determines the location of the primary binding event, DNA topology emerges as a key factor shaping how the complex organizes and stabilizes once binding has occurred. In this view, positive supercoiling act as an allosteric modulator of the conformational landscape accessible to the protein–DNA complex, while negatively supercoiled supports broader rearrangements and a less constrained configurational landscape. By restricting conformational heterogeneity and stabilizing ordered binding ensembles, positive supercoiling biases the system toward a subset of structurally and dynamically distinct states. This effect is evident both in the reduced plasticity of the DNA-binding region and in the more focused redistribution of additional protein dimers along the same DNA template.

These results suggest that topology does not only modulate whether binding occurs, but also how the bound state is organized and how alternative protein binding are distributed along the DNA. The local hydrogen-bond and DNA compaction readouts support the same general view, namely that the immediate protein–DNA interface reflects the broader topological bias of the substrate. These local observations were then placed in a broader structural context through coarse-grained simulations of the plasmid. The plasmid analysis revealed that positively supercoiled DNA occupies a narrower and more compact conformational basin, whereas negatively supercoiled DNA samples a broader and more heterogeneous landscape. Importantly, the same sign-dependent ordering is preserved in the matched 336-bp segment and in the corresponding mcDNA systems, supporting the use of the minicircle as a mechanistically informative local proxy of the plasmid response rather than as an isolated model. In this sense, the multiscale framework developed here connects plasmid-scale topology with local structural readouts and provides a coherent way to interpret how global DNA mechanics are translated into protein conformational behavior.

Taken together, our findings support a model in which local protein–DNA recognition and global DNA topology are tightly coupled. Positive supercoiling, which remains comparatively less explored than negative supercoiling in this context, emerges here as a structurally distinctive state capable of promoting a more compact DNA ensemble and a more ordered protein-bound configuration. More broadly, this work identifies DNA supercoiling as an active regulator of transcription factor recognition and suggests that topology-dependent tuning of protein binding may represent a general layer of regulation for bZIP proteins and, potentially, for other DNA-binding factors whose activity depends on DNA shape, flexibility, and torsional state. These observations suggest that DNA topology can encode regulatory information orthogonal to sequence, contributing to the functional outcome of protein–DNA interactions without altering the underlying nucleotide code. In this sense, local supercoiling may be interpreted as a form of topological code that modulates binding stability, conformational selection, and higher-order organization of protein assemblies on DNA. Such a mechanism provides a plausible framework for integrating mechanical stress, local DNA geometry, and protein conformational dynamics *in vivo*, particularly in contexts where positive supercoiling is transiently generated by active transcription or constrained chromosomal architecture.

## 4. Experimental section

### GCN4 expression and purification

The GCN4 plasmid pGCNK58 for *E. coli* expression was kindly provided by Prof. A. Palmer without any tag (24). The protein was expressed and purified in *E. coli* BL21 (DE3) cells, which were transformed and grown in 4 L of Luria Broth media at 37 °C until reaching an optical density (OD600) of 0.5. Protein expression was induced with 1 mM IPTG and allowed to proceed for approximately 4 hrs. Lysis was achieved by sonication in 25 mM HEPES and 50 mM NaCl, pH 7.5.

The protein was purified by Ion Exchange Chromatography on AKTA Go system with Cityva HiTrap SP FF 5 ml column. Elution was performed by NaCl gradient from 50mM to 1M. Fractions were checked by SDS-PAGE, then pooled, concentrated to 1.2 mM, and dialyzed in 25 mM HEPES and 50 mM NaCl. Independently, an attempt was also carried out using a further step of purification by HPLC according to the available protocol. Purity was checked by SDS-PAGE and MALDI TOF Mass Spectrometry. This technique was performed using the advanced platform of the Institut de Biologie Structurale of the EPN campus in Grenoble. The final purity was estimated to be >98%. Protein concentrations, determined by UV spectroscopy, were defined with respect to the monomer.

### Preparation of plasmid DNA in distinct topological states

The commercial plasmid pUC19 (2686 bp) was isolated from E. coli using the PureLink HiPure Midiprep kit (Thermo Fisher Scientific). This preparation, obtained in the negatively supercoiled form pDNA(SC-), served as the substrate for all subsequent topological conversions. Nicked open-circular DNA was generated by incubating pDNA(SC-) (1 µg) with 1 U mg-1 of the strand-specific nicking endonuclease Nt.BspQI (New England Biolabs) in 1 x Nt.BspQI buffer at 50° C for 2 h. Linear DNA was produced by complete digestion of pDNA(SC-) with the single-site restriction enzyme EcoRI (New England Biolabs) under saturating enzyme conditions in the manufacturer’s buffer at 37 °C for 2 h. Positively supercoiled pDNA(SC+) was obtained by treating pDNA(SC-) (30 µg) with 300 µg ml-1 reverse gyrase from Saccharolobus solfataricus (24) in 35 mM Tris-HCl (pH 7.0), 0.1 mM EDTA, 30 mM MgCl2, 2 mM DTT and 1 mM ATP. Reactions were incubated at 90 °C for 5 min and stopped with 2% SDS (7). After each enzymatic step, DNA was purified by organic extraction and ethanol precipitation. Samples were extracted once with buffer-saturated phenol and twice with phenol:chloroform:isoamyl alcohol (25:24:1, v/v), precipitated with 0.1 vol 3 M sodium acetate and 3 vol cold absolute ethanol, incubated at –20 °C overnight, pelleted by centrifugation, air-dried, and resuspended in nuclease-free water.

### Electrophoretic mobility shift assay (EMSA)

Plasmid DNA binding assays were performed in 10 µL reactions volumes containing 20 mM Tris–HCl (pH 8.0), 50 mM KCl, 0.1 mM DTT and 10 % (v/v) glycerol, in which pDNA (3 nM) in four different topological states (negatively or positively supercoiled, open-circular, linear) was incubated with GCN4 with increasing concentrations of GCN4 (5– 1000 nM) for 10 min at 37 °C. Samples were loaded directly onto a 1.2 % agarose gel prepared in 0.5 × TBE buffer and run at 100 V for 2 h at room temperature. Gels were stained with ethidium bromide (1 µg mL^-1^), rinsed in deionized water and visualized under UV illumination with a VersaDoc 4000 imaging system (Bio-Rad). Band intensities were quantified using Quantity One software (Bio-Rad) and the fraction bound-pDNA(GCN4-pDNA Complexes) was quantified, as previously described (29). Binding curves were generated by plotting the fraction of bound DNA as a function of protein concentration and fitted using a one-site specific binding model to determine the dissociation constant (k_d_), using GraphPad Prism (GraphPad Software).

### BLI assays

OctetRED96TM (ForteBio) was used to immobilize biotinylated oligonucleotides by streptavidin-coated biosensors (Octet^®^ High Precision Streptavidin Biosensors, Sartorius). Samples and reaction buffers 1X (Kinetic Buffer, Sartorius) were in black 96-well plates (OptiPlate-96 Black, Black Opaque 96-well Microplate, PerkinElmer) in a reaction volume of 200 µL per well with 1000 rpm shaking for each step. Before starting the experiment, the biosensors were hydrated for 15 min in 200 µL of pure water. Baseline was recorded for 60 s, after which the biosensors were immersed in wells containing a fixed concentration of biotinylated double-stranded DNA oligonucleotide (bio-oligo, 0.1 µM) during the loading step (600 s). Following a washing step in the kinetic buffer (180 s), a quenching step in biocytin (300s) and another washing step to remove the excess of blocking molecule (180 s), the Association of increasing amounts of GCN4 protein (0-500 nM) was recorded for 300 s, by immersing the coated sensor in wells containing the protein diluted in kinetic buffer 1X, as well as the dissociation step was recorded by immersing the sensor tips in wells containing pure kinetic buffer 1X (600 s). The sensorgram was analyzed using Global 1:1 fitting of association and dissociation curves with the Data Analysis 9.0 software. All the BLI experiments were performed at 30 °C. The competition experiments were performed using the same conditions. A fixed concentration of GCN4 (0.1 µM) was loaded onto pre-immobilized bio-oligos for 600 s (association step), followed by washing and quenching steps. Subsequently, the bio-oligo/GCN4 complex was immersed in different wells (dissociation step, 600 s) containing plasmid DNA in various amounts (15.6 ng to 250 ng) as a competitor. The sensorgram was analyzed using Local 1:1 fitting of dissociation curves with Data Analysis 9.0 software. The experiment was performed similarly for all the DNA topologies, at 30 °C with 1000 rpm shaking, with one well containing only buffer (no DNA competitor) being used as a control.

### Circular-dichroism spectroscopy

Circular-dichroism (CD) spectra were collected on a J-1500 spectropolarimeter (JASCO, Tokyo, Japan) fitted with a 1 mm quartz cuvette (Hellma), a Peltier-controlled cell holder and a Julabo F-25 circulator maintained at 25 °C. Spectra were scanned from 190 to 300 nm at 100 nm min^-1^ with a 0.5 nm bandwidth; three accumulations were averaged for each sample and the corresponding buffer baseline was subtracted automatically. The following samples were examined: free GCN4 (5 µM); free plasmid DNA (16 nM) in four topological states (negatively supercoiled, positively supercoiled, open-circular, linear); and GCN4–DNA complexes prepared at a 1:300 protein: DNA molar ratio. All mixtures were incubated for 5 min at 25 °C before data acquisition. Effects of GCN4 on DNA conformation were assessed in the 260–300 nm window, where protein contributions are negligible because the GCN4 leucine-zipper contains only a single aromatic residue. Raw ellipticity is reported in millidegrees (θ, mdeg); curves were smoothed with standard noise-reduction algorithms provided by Prism (GraphPad).

### Computational methods

#### Coarse-grained and atomistic MD simulations

Coarse-grained simulations of the 2695-bp plasmids were used to describe topology-dependent global DNA organization, whereas a reduced 336-bp minicircle was used for local, site-resolved atomistic analyses. The minicircle was centered on the plasmid region containing the specific AP-1 sequence landmarks considered throughout the study. Two DNA states were considered for the 2695-bp DNA plasmid model system, pDNA(SC-) and pDNA(SC+), whereas four local DNA states were considered for the 336-bp model system: mcDNA(SC+), mcDNA(SC-), mcDNA(LIN), and mcDNA(NICK), where mcDNA(NICK) denotes the nicked open-circular control and mcDNA(LIN) the corresponding linearized DNA with free ends. Representative back-mapped configurations of the control mcDNA(NICK) and mcDNA(LIN) systems are shown in Supplementary Figure S4.

The 336-bp minicircle models were generated using NAB/AmberTools from constructs containing the specific AP-1 sequence binding site within the selected plasmid region (41). The plasmid topologies were prepared at the target supercoiling densities pDNA(SC-) (σ = −0.156) and pDNA(SC+) (σ = +0.156) using native oxDNA topology-generation tools (42,43), whereas the minicircle states mcDNA(SC+) and mcDNA(SC-) were prepared at the corresponding local supercoiling densities, and relaxed control states were generated as nicked open-circular and linearized DNA. Initial GCN4-DNA complexes were generated from the crystallographic GCN4-DNA complex (PDB 1YSA), following the same general strategy adopted in our previous collaborative work, by placing one dimer at the AP-1 sequence site and four additional dimers at non-specific positions elsewhere on the minicircle. In all cases, the protein-binding sites were separated by at least 35 bp to minimize trivial steric overlap and to probe topology-dependent redistribution beyond the primary binding event. Geometry building, circularization, sequence mapping, docking preparation, and backmapping were carried out with NAB/AmberTools, oxView, and custom Python scripts (41,42).

DNA topology of the plasmids was characterized using oxDNA2, which provides a nucleotide-level coarse-grained description suitable for capturing supercoiling-dependent changes in DNA structure, mechanics, and mesoscale conformational organization (43). Simulations were performed at 27 °C and 0.15 M salt concentration using average-sequence interaction parameters. Initial coarse-grained structures were first subjected to a short Monte Carlo relaxation to remove local steric stress, followed by a molecular-dynamics relaxation stage. Production trajectories were then propagated in the NVT ensemble using dt = 0.003, the Bussi thermostat (bussi_tau = 1000, newtonian_steps = 53), and no external biasing forces. One trajectory was generated for each plasmid topological state. Each system was simulated for 1000 ns after equilibration, and the first 30% of each trajectory was excluded from structural averaging to focus on the quasi-stationary regime.

Atomistic simulations focused on the four minicircle systems mcDNA(SC+), mcDNA(SC-), mcDNA(LIN), and mcDNA(NICK), which explicitly resolve local DNA deformation and protein-DNA contacts in the plasmid region containing the specific binding site and the additional AP-1/AP-1-like sequence landmarks. DNA was described with the parmbsc1 force field (44), the protein with the Amber99 force field, and water molecules with the TIP3P model. Each system was simulated for 100 ns after equilibration, and the first 30% of each trajectory was excluded from structural averaging to focus on the quasi-stationary regime. Where exact low-level production settings were not recoverable from the available local files, no unsupported values were inferred and only robustly documented simulation details were retained in the present version.

#### Analysis of simulations

Trajectory analysis was organized at complementary structural levels. Protein-centered readouts included residue-wise flexibility, secondary-structure persistence, overall helical content, and two geometric descriptors, Dih1 and Dih2, used to monitor the relative rearrangement of the two DNA-binding basic regions and the leucine-zipper dimerization core, respectively. Protein flexibility was estimated from root-mean-square fluctuations (RMSF) of Cα atoms along the sequence, thereby reporting residue-resolved positional fluctuations in the protein backbone. Secondary structure was assigned with DSSP (45), and the overall helical content (%) of the bZIP domain was monitored throughout the trajectories to evaluate topology-dependent stabilization of the bound protein conformations. Residue fluctuations and RMSF profiles were obtained from GROMACS-based analyses (46).

For cross-scale comparison, the 336-bp local sequence used in the minicircle was first identified unambiguously within the plasmid by exact sequence matching on both strands. The corresponding coarse-grained plasmid segment was then extracted frame by frame from the plasmid trajectories. Because DNA supercoiling is formally defined for topologically closed molecules, no independent topological state was assigned to the extracted 336-bp segment; instead, that segment was treated as a local structural reporter embedded within a globally supercoiled plasmid. Representative coarse-grained plasmid configurations and the matched 336-bp segment were subsequently backmapped to atomistic resolution and relaxed using the same DNA force field adopted for the atomistic simulations (parmbsc1) (44). This procedure ensured that compactness-related descriptors, including DNA radius of gyration, were evaluated only after conversion of the coarse-grained-derived structures to a common atomistic representation followed by relaxation with the parmbsc1 force field. In the same spirit, the free-energy landscapes discussed in the cross-scale comparison were interpreted on the basis of the corresponding atomistically relaxed representations, so that the qualitative conformational ordering could be compared on a consistent structural level between coarse-grained-derived and fully atomistic models.

DNA radius of gyration was computed on DNA-only atom selections after atomistic backmapping and relaxation of the coarse-grained-derived structures. For the cross-scale comparison, DNA compactness was therefore evaluated on a common atomistically relaxed representation in three matched contexts: representative plasmid conformations derived from the coarse-grained trajectories, the extracted 336-bp segment from the plasmid coarse-grained trajectories, and the atomistically simulated 336-bp minicircle. This comparison was interpreted qualitatively, as a test of whether the same sequence window preserves the same compaction ordering across models, rather than as an attempt to equate absolute coarse-grained and atomistic Rg values.

To quantify sequence-resolved recognition, the 336-bp reference was explicitly annotated for the AP-1/AP-1-like motifs considered in this study. Custom Python workflows were then used to monitor, frame by frame and dimer by dimer, sequence distance, spatial distance, and motif overlap relative to these landmarks. This analysis was used to identify the dominant specifically engaged dimer, quantify redistribution of non-primary dimers across the four DNA states, and distinguish persistent docking from transient sampling of secondary AP-1/AP-1-like regions.

Conformational heterogeneity was characterized in two complementary ways. First, coarse-grained trajectories were backmapped to the atomistic level and then projected onto backbone torsional descriptors and analyzed by time-lagged independent component analysis (tICA) to identify the dominant slow conformational modes of the coarse-grained trajectories (47). Representative coarse-grained conformations were further identified by k-medoids clustering in the reduced tICA space, with the number of clusters selected by silhouette analysis (48). These representative states were used for structural inspection and for the cross-scale comparison between the plasmid, the matched 336-bp plasmid segment, and the 336-bp minicircle model. For cross-scale comparison, the corresponding representative coarse-grained conformations were backmapped and atomistically relaxed before qualitative comparison of the resulting free-energy organization. Second, atomistic conformations were classified using physically motivated local features, including local DNA curvature, local DNA compactness, and dimer-position metrics relative to the annotated AP-1/AP-1-like motifs. Five candidate basins were initially generated; after robustness screening, two representative basins (states A and B) were retained for each system, and up to 20 representative frames per basin were extracted for structural inspection and energetic analysis.

#### Binding energetics analysis

Binding free energies for the DNA-protein complexes were estimated using the Molecular Mechanics Generalized Born Surface Area (MM/GBSA) method, which combines molecular mechanics energy terms with an implicit-solvent treatment based on the Generalized Born formalism (49). Calculations were carried out with gmx_MMPBSA interfaced with AmberTools/Amber22 (50). Although MM/GBSA does not provide absolute binding free energies with high precision, it offers an internally consistent comparative framework for evaluating relative energetic trends across DNA topologies, dimers, and conformational states.

In the MM/GBSA framework, the binding free energy is expressed as

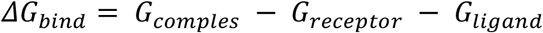

with each free-energy term decomposed as

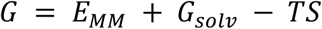

where the total free energy includes the molecular mechanics contribution and the polar and nonpolar solvation terms (49).

In the single-trajectory approximation, receptor and ligand configurations are extracted from the complex trajectory; under this assumption, internal bonded terms cancel formally between bound and unbound states, so the comparison is dominated primarily by van der Waals, electrostatic, and solvation contributions (49). Because no entropic term was explicitly included here, the reported MM/GBSA values should be interpreted as effective comparative binding energies rather than absolute free energies (49).

Two complementary protocols were applied: (i) long-window calculations on the final 30 ns of the reference trajectories for the systems in which this segment was explicitly prepared, and (ii) state-resolved calculations on two representative conformational basins for all four minicircle systems. In the long-window protocol, all saved frames from the final 30 ns were analyzed, corresponding to 600 frames at 50-ps stride. In the state-resolved protocol, all five GCN4 dimers were analyzed independently, allowing topology-dependent redistribution of stabilization to be assessed at dimer resolution.

A single-trajectory MM/GBSA protocol was adopted as a simple and internally consistent comparative framework. Standard Generalized Born settings were used, with (igb = 5) and a physiological salt concentration of 0.15 M (49). No entropic correction was included. Accordingly, MM/GBSA was used here as a comparative energetic descriptor rather than as an absolute predictor of binding affinity, and interpretation was restricted to within-study trends across systems, dimers, and conformational states (49).

## Acknowledgements

We wish to express our deepest gratitude to Prof. Giuseppe Perugino (Pino), whose insightful scientific discussions and guidance have been invaluable. His passion and clarity of thought continue to serve as an inspiration. This work is dedicated to his memory.

We thank Jonathan Leon Moro for cloning the pUC19-Arg plasmid, used as a control in experiments not shown.

Part of the instrumentation used in this study was funded through: IR0000032 ITINERIS (Italian Integrated Environmental Research Infrastructures System (CUP B53C22002150006) under the EU Next Generation EU PNRR programme); SEE LIFE-StrEngthEning the ItaLIan InFrastructure of Euro-bioimaging (Area ESFRI “Health and Food”-IR0000023); the National Academic Infrastructure for Supercomputing in Sweden (NAISS), partially funded by the Swedish Research Council through grant agreement no. 2022-06725.

