## Supplementary material for "DNA supercoiling modulates bZIP transcription factor–DNA interaction"

**Figure S1.** *Dose-response curve of GCN4/bioOligo interactions.* Association and dissociation curves of various concentrations of GCN4 (0-500 nM) after the interaction with a fixed concentration (0.1 μM) of pre-immobilised biotin-oligonucleotide.

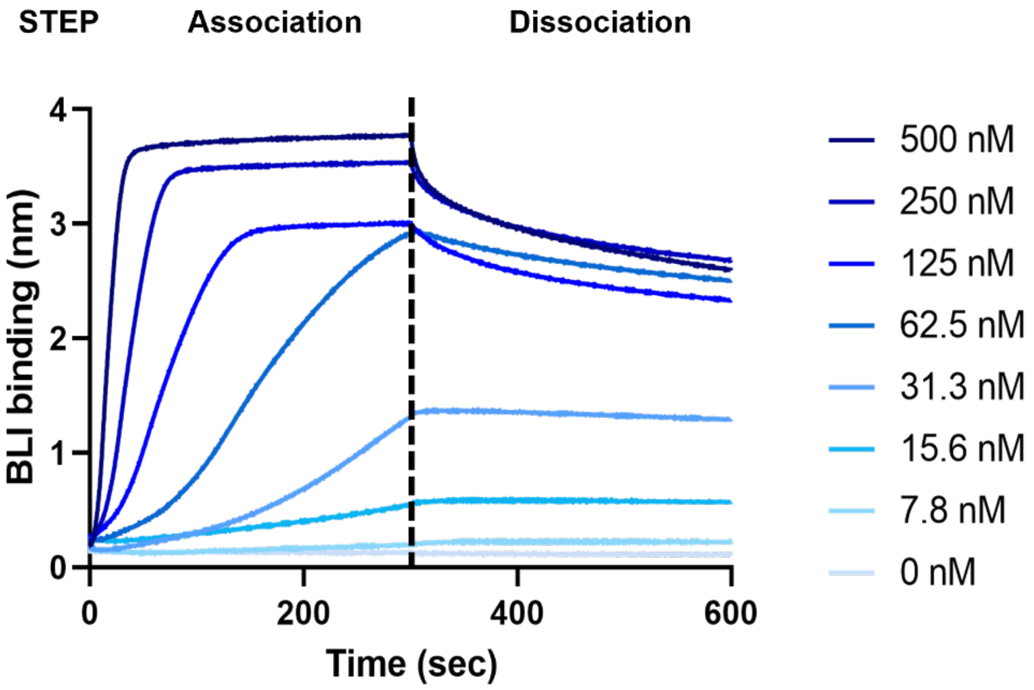

**Figure S2.** Snapshots of mcDNA(SC-) and of the corresponding complex with GCN4. A) The input coarse-grain in open-circle configuration; B) a representative snapshot (380 ns) from the coarse-grain simulation; C) docked dimeric GCN4 used for coarse-grain equilibration; D) the atomistic configuration used as input for atomistic simulations obtained from back-mapping of the coarse-grain structure.

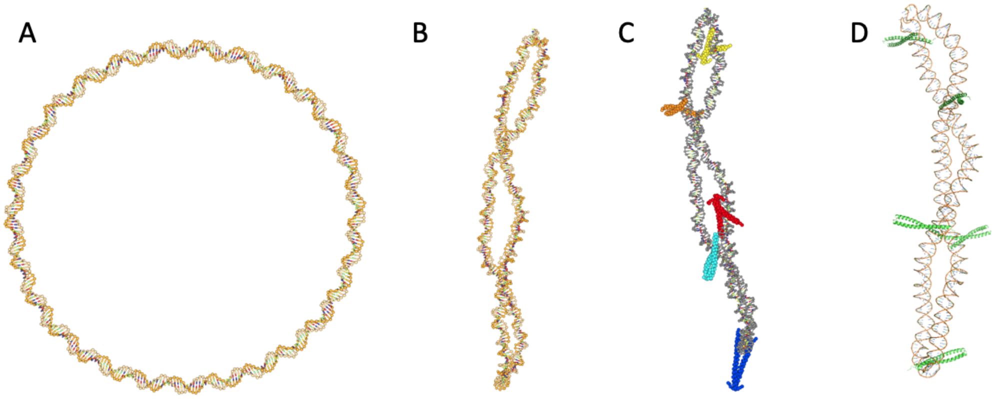


**Figure S3.** Snapshots of mcDNA(SC+) and of the corresponding complex with GCN4. A) The input coarse-grain in open-circle configuration; B) a representative snapshot (430 ns) from the coarse-grain simulation; C) docked dimeric GCN4 used for coarse-grain equilibration; D) the atomistic configuration used as input for atomistic simulations obtained from back-mapping of the coarse-grain structure.

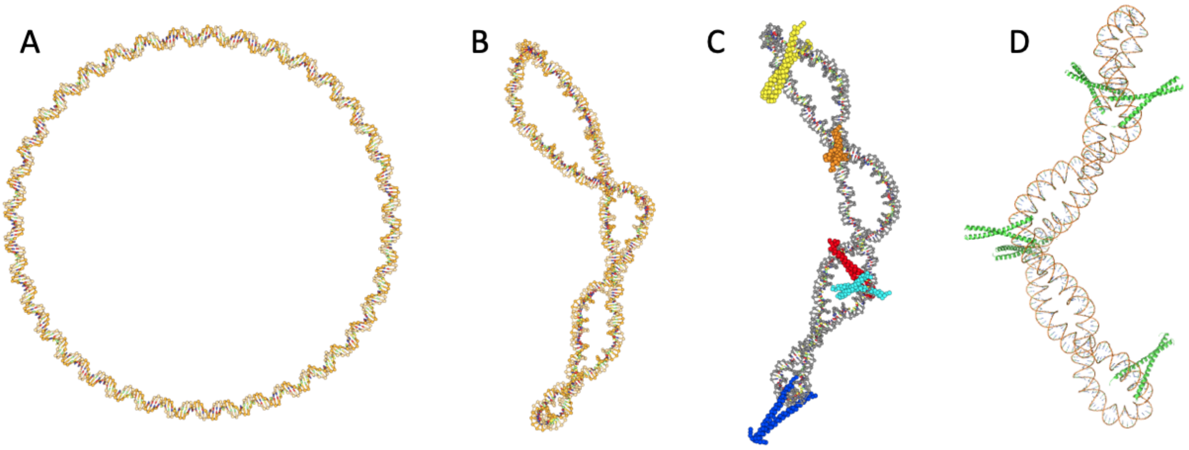

**Figure S4.** Snapshots of the back-mapped configurations at the end of the simulations of mcDNA(NICK) and mcDNA(LIN). The nicked sites are highlighted with an asterisk.

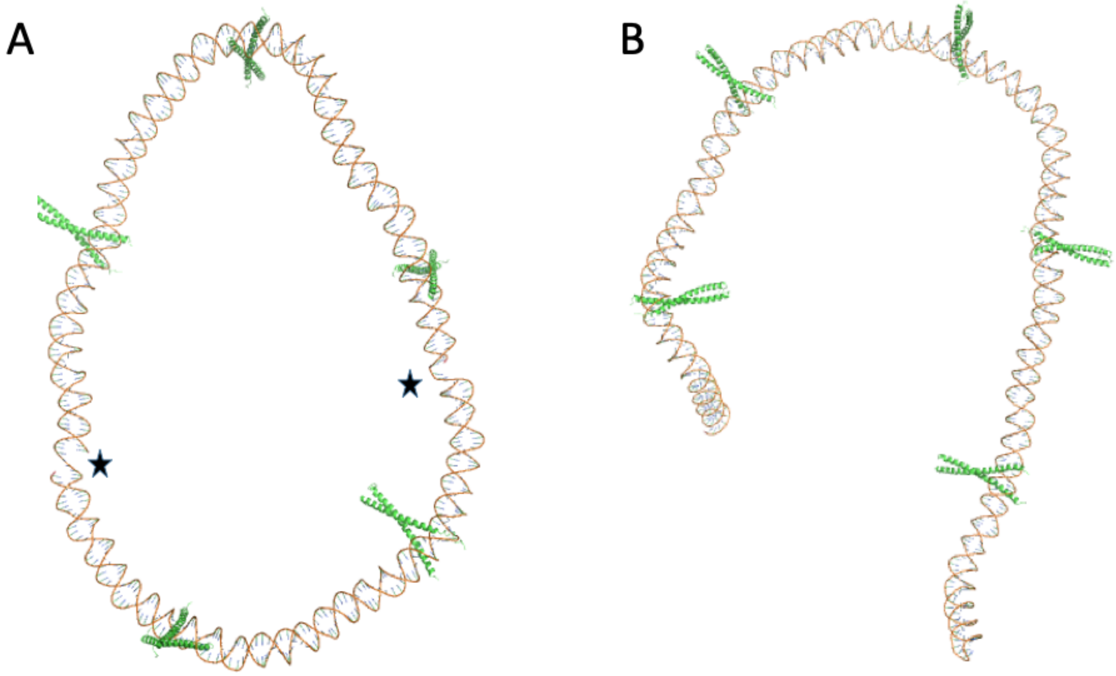

**Figure S5.** Conformational dynamics of GCN4 functional domains. (A) Schematic of the GCN4 dimer bound to DNA (modified from PDB 1YSA) showing the two geometric descriptors used in the trajectory analysis: Dih1 across the two DNA-binding basic regions and Dih2 across the two leucine-zipper helices. (B) Dih1/Dih2 projection sampled along the trajectories, showing topology-dependent redistribution of the DNA-binding arms while the zipper core remains comparatively constrained. (C) Representative conformers illustrating the broader basic-region plasticity in mcDNA(SC-) and the more compact ensemble sampled in mcDNA(SC+).

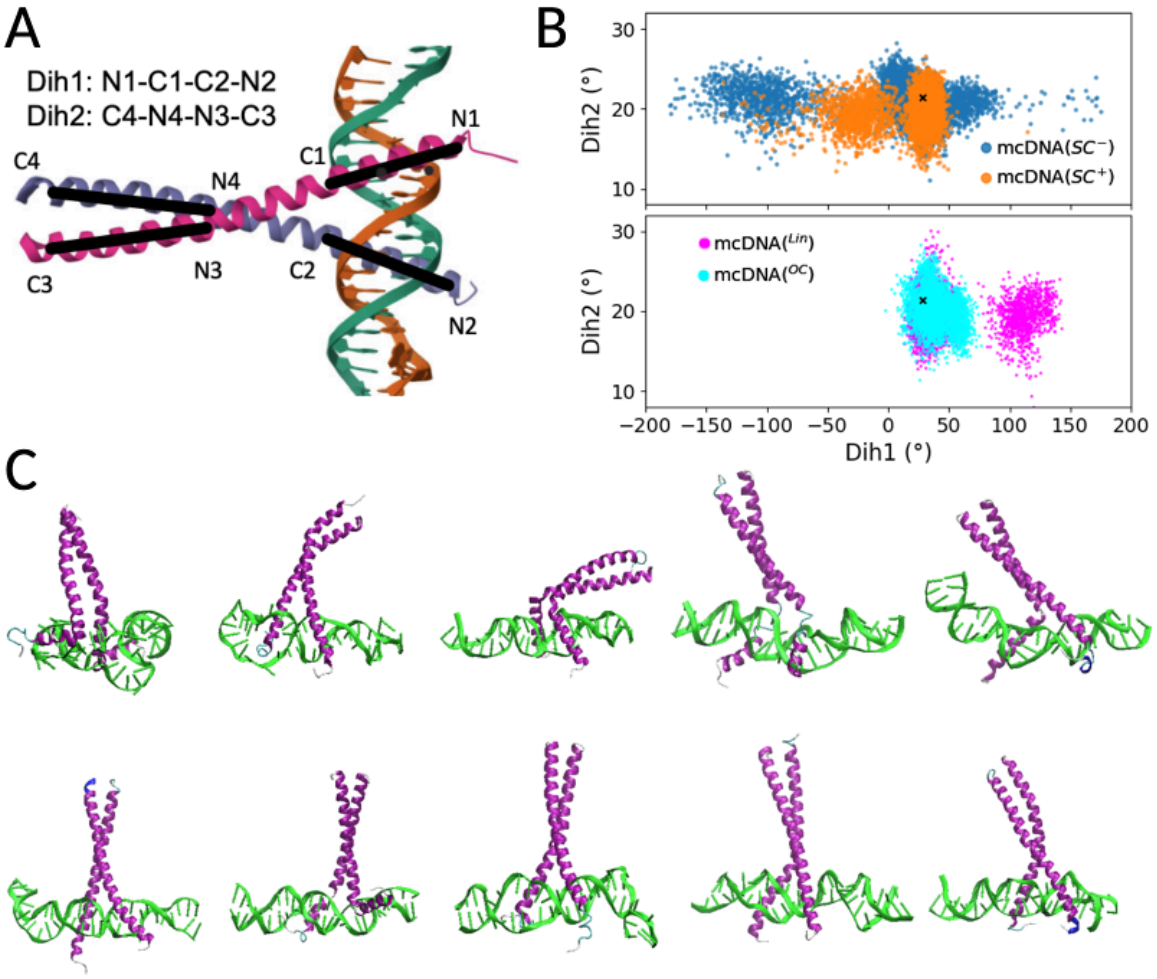

**Figure S6**. Cross-scale radius-of-gyration validation of the topology-dependent compaction bias. The figure compares the ordering of DNA compactness across three matched apo contexts: plasmid coarse-grained DNA, the matched 336-bp coarse-grained segment extracted from the plasmid, and the apo atomistic 336-bp mcDNA. The plasmid radius of gyration is retained as the global reference.

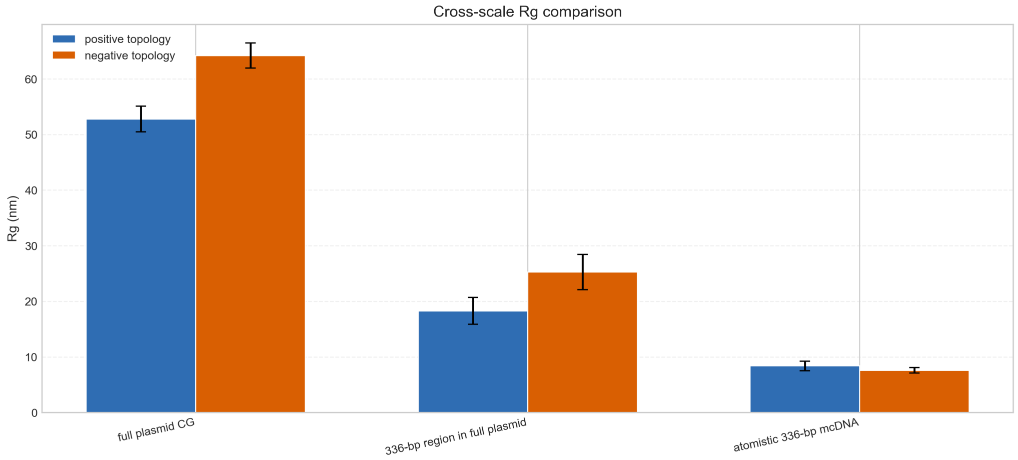


**Figure S7.** Protein-DNA hydrogen bonds and local DNA radius of gyration across the atomistic mcDNA systems.

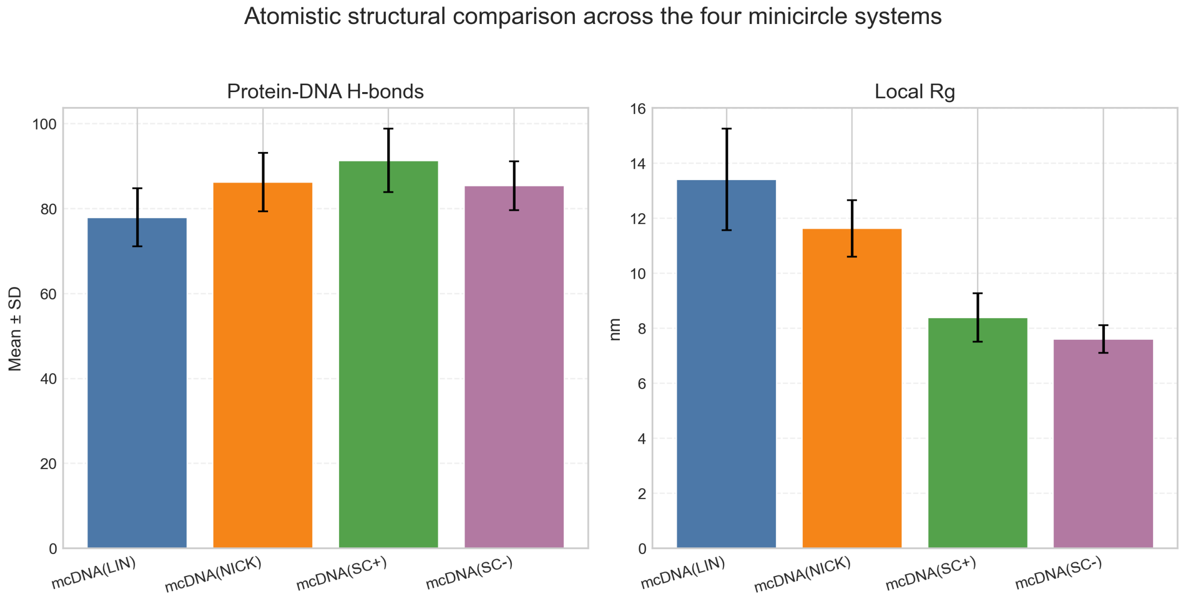

**Figure S8**. Residue-resolved secondary-structure persistence maps for the two helices of the GCN4 basic region in mcDNA(SC-) and mcDNA(SC+) complexes. The heatmaps report the time evolution of DSSP-assigned structural states for helix 1 and helix 2, highlighting the broader secondary-structure heterogeneity retained in mcDNA(SC-) and the more persistent helical organization sampled in mcDNA(SC+).

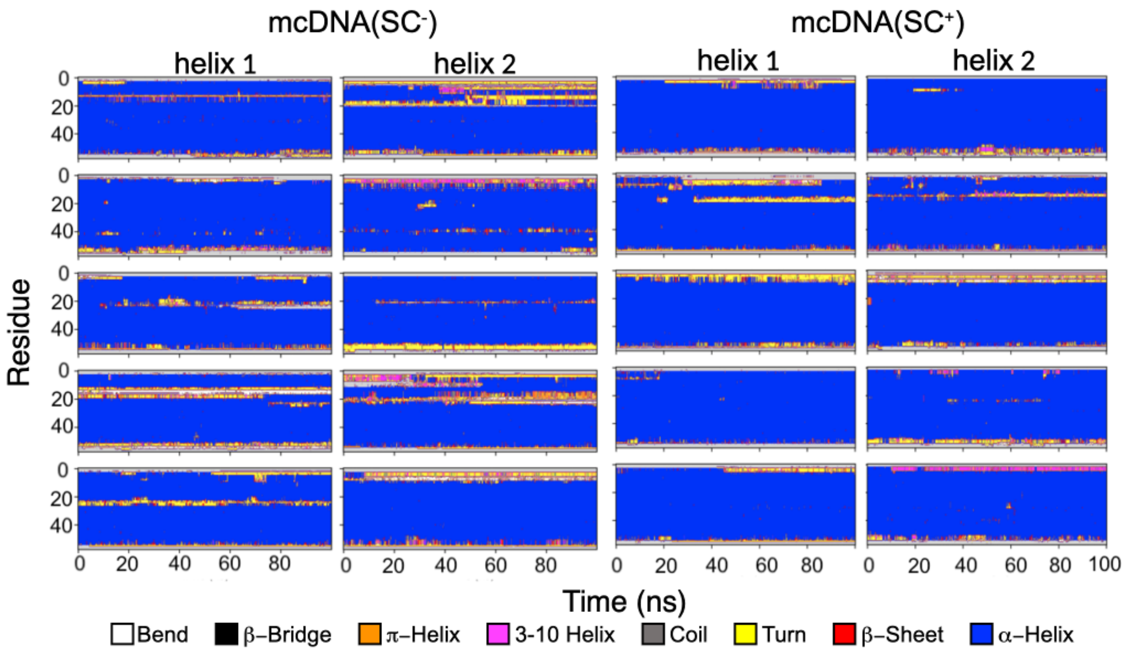

**Figure S9**. Per-dimer RMSF and helicity profiles across the mcDNA(SC-) and mcDNA(SC+) complexes. For each of the five GCN4 dimers, the residue-wise RMSF (red) and helical content (blue) are reported to illustrate topology-dependent differences in conformational plasticity of the DNA-binding region.

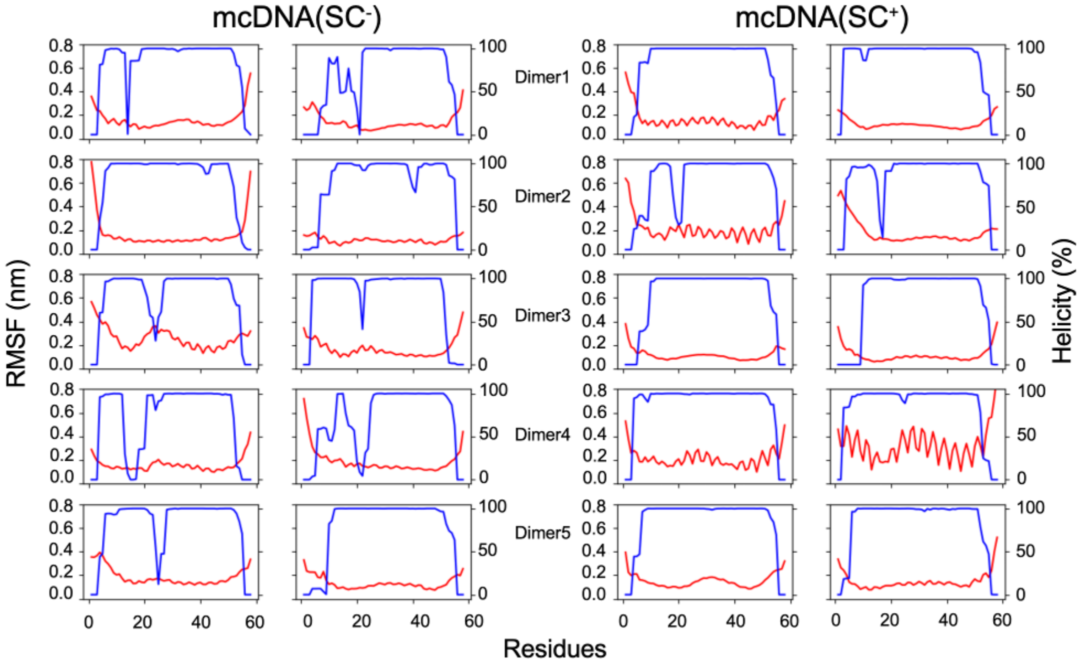


**Figure S10**. Ranking by best non-primary AP-1/AP-1-like approach across the four mcDNA systems. The plot reports the mean minimum approach distance of the best non-primary dimer to AP-1/AP-1-like motifs, highlighting the more focused secondary organization observed in mcDNA(SC+).

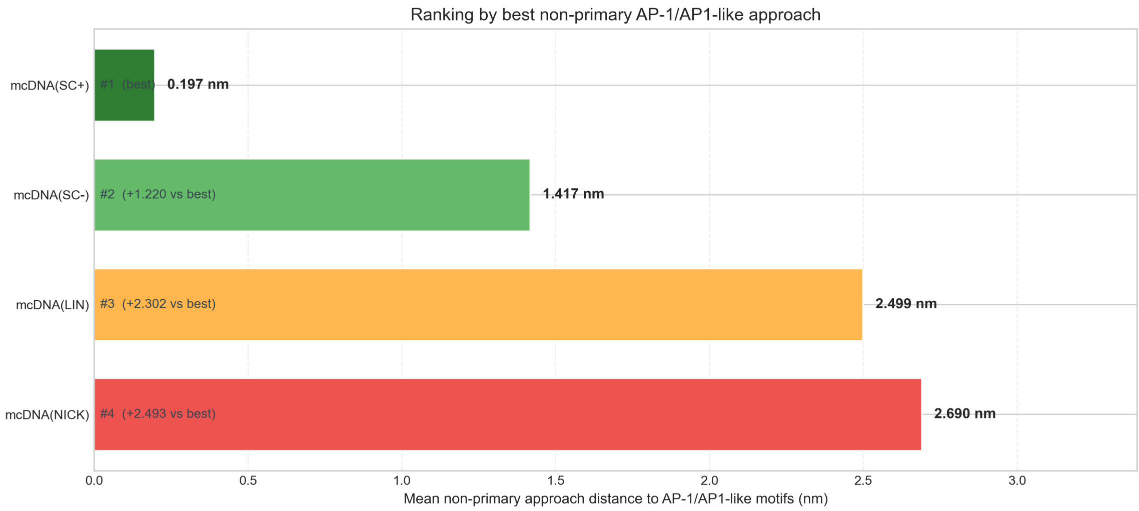

**Figure S11.** Comparative MM/GBSA mean values across the five dimers in the four mcDNA systems. Bars report the within-study average effective binding-energy estimates used only as supportive energetic descriptors, consistent with the topology-dependent trends discussed in the main text.

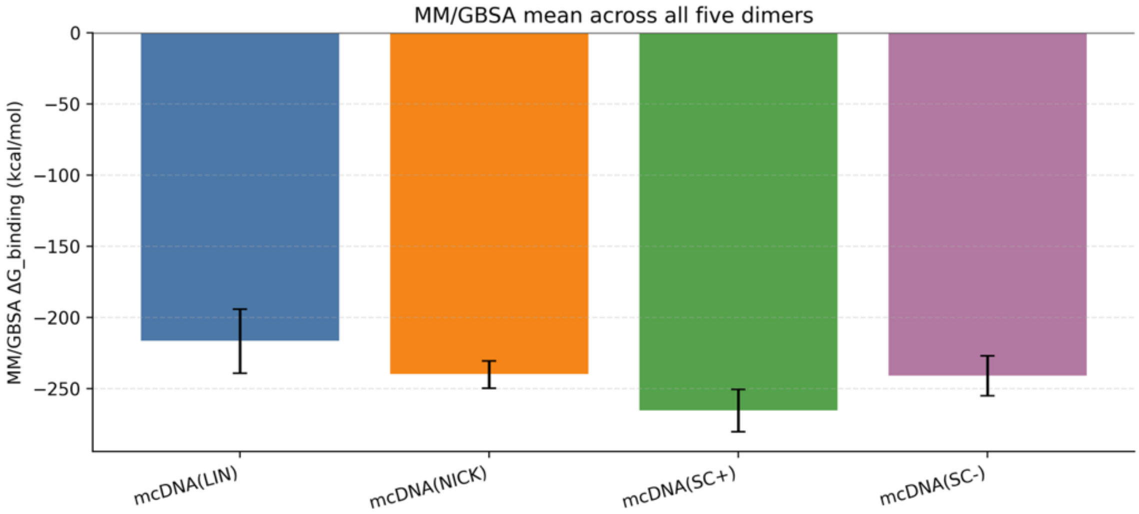


**Figure S12.** Snapshots of -pDNA(SC+): A) The input CG in "open-circle" configuration after relax and MD relax steps using oxDNA software package; B) A representative snapshot from the CG simulation after some MD relax steps.

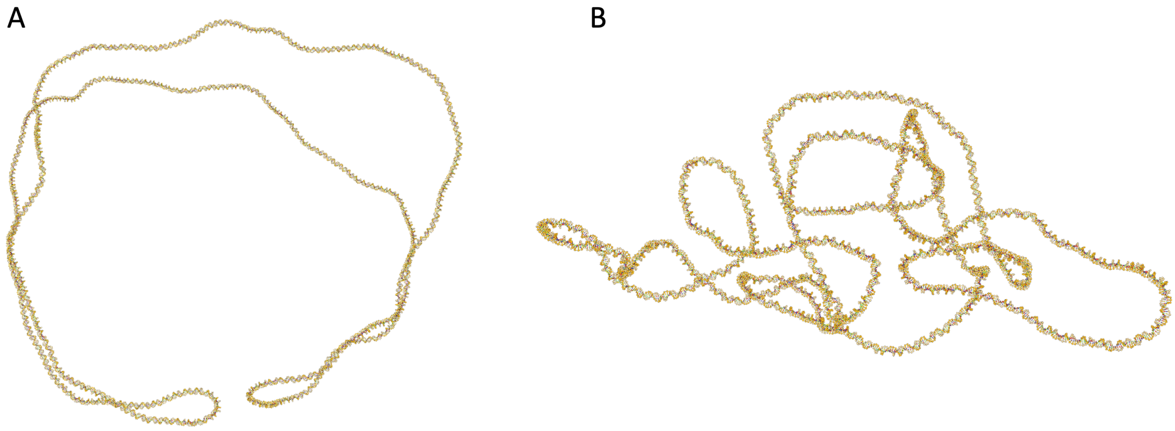
